# Multi-omics integration at cell type resolution uncovers gene-metabolite mechanisms underlying osteoarthritis heterogeneity

**DOI:** 10.1101/2025.09.26.678886

**Authors:** Abhishek Ojha, Paramita Chatterjee, Linda E. Kippner, Hazel Y. Stevens, Scott D. Boden, Hicham Drissi, Kenneth Mautner, Carolyn Yeago, Krishnendu Roy, Saurabh Sinha

## Abstract

Metabolic dysregulation is an important factor for osteoarthritis pathogenesis, but comprehensive studies of underlying mechanisms and pathways are rare. We analyzed newly generated metabolomics data on bone marrow from 119 osteoarthritis patients, along with single-cell transcriptomics data to reconstruct networks of gene-metabolite associations at cell type resolution. Hubs of these networks – cell type-specific as well as pan-cell type hubs – revealed key molecular factors of osteoarthritis heterogeneity. Systems-level analysis of hubs revealed major roles for glycerophospholipid, glycerolipid and sphingolipid metabolism pathways, along with lipid signaling. We used Machine Learning models of gene-metabolite relationships to discover cell types most relevant to each metabolite. Integrative analysis of disease severity scores along with multi-omics data revealed a shift in specific immune cell subtypes in low versus high grade disease. We conclude that leveraging gene-metabolite covariation in a patient cohort can uncover underlying molecular mechanisms, overcoming the challenges posed by high dimensionality of multi-omics data.

## INTRODUCTION

Osteoarthritis (OA) is the most common type of arthritis, which primarily affects older adults and is characterized by the gradual breakdown of joint tissues, leading to pain, stiffness, and reduced mobility. Metabolites play a crucial role in the development and progression of OA [1] [2] making metabolomic studies a promising approach to understanding OA etiology. Such studies have shown alterations in metabolite concentrations to aid diagnosis and prognosis of OA [1, 3] and have identified OA-associated metabolic pathways [4]. Transcriptomics technologies have also been employed to probe the molecular basis of OA, identifying gene expression correlates of OA progression [5, 6] with potential applications to disease monitoring and therapeutics [7].

While metabolomics- and transcriptomics-based strategies have individually shown promise, integrative approaches that provide a holistic view of these complementary omics profiles in OA remain largely unexplored. Cell metabolism and gene expression are intertwined processes whose participants (metabolites and proteins) come together through diverse pathways to shape phenotypic outcomes [8]. Thus, their respective omics profiles can fill in contextual gaps in each other and if analyzed jointly can reveal a more precise mechanistic characterization of phenotypes than either metabolomics or transcriptomics can by itself [9]. For instance, Xie et al. [10] relied on RNA-metabolite covariation analysis to evaluate hypotheses regarding metabolic impacts of cancer and its progression, while Hu et al. [11] analyzed the two “omes” at spatial single-cell resolution to discover unique metabolic states within cell types in lung cancer. The value of such joint analysis has been convincingly demonstrated by multiple pan-cancer studies, revealing mechanistic interactions as well as metabolic remodeling [12, 13]. However, the promise of multi-omics synergy is yet to be realized in the context of OA.

Any integrative analysis that aims to jointly study transcript and metabolite variation in complex tissues must contend with the confounding effects of cell type heterogeneity. Gene expression programs are well-documented to vary across cell types [14] and a growing body of work highlights similar variations in metabolic profiles [15]. Thus, prior studies examining gene-metabolite covariation, which are mostly restricted to “bulk” omics profiles, are likely to have overlooked critical cell type-specific relationships. In contrast, a multi-omics covariation analysis leveraging cell type-resolved data holds greater potential for uncovering these relationships.

Multi-omics analysis of a disease cohort presents the most daunting challenges of data science – sparsity and high dimensionality. One might consider training a predictive model relating the phenotype of subjects (e.g., case/control or disease severity) to their multi-omics profiles, followed by model interpretation [2] [16]. However, in practice a patient cohort with multi-omics profiles rarely has more than hundreds of patients, while the number of omics features is in the thousands. This makes it challenging to train models and extract mechanistic insights from them. Here, we explore a pragmatic solution to this methodological challenge. Rather than train “omics-to-phenotype” models, the strategy is to systematically examine covariations between genes and metabolites profiled in disease-relevant tissue from a cohort of patients with varying disease severity. This yields a cross-omics covariation network whose “hubs” (molecules that covary with many other molecules) can point us to key factors related to the phenotype. This strategy has been successfully used in many transcriptomics studies, where hubs of co-expression networks point to important regulators of the phenotype of interest [17].

Our work presents a comprehensive multi-omics analysis of OA patient data at cell type resolution (**Figure 1A**). We generated liquid chromatography-mass spectrometry (LC-MS) profiles from bone marrow aspirate concentrate samples of ∼120 OA patients who participated in the MILES clinical trial [18], and analyzed these jointly with single-cell whole-transcriptome (scRNA-seq) profiles of the same samples, using rigorous statistical techniques and machine learning tools. We constructed maps of statistically significant gene-metabolite relationships for each of 17 cell types and identified cell type-specific as well as pan-cell type hub genes and metabolites, highlighting their roles in inflammatory processes, cartilage homeostasis, and metabolic dysregulation. Focusing on gene-metabolite pairs with prior evidence of pathway proximity (involved in the same or successive reactions) revealed phosphatidylinositol, phosphatidylcholine, sphingolipid and glycerophospholipid metabolism, along with lipid signaling, as critical pathways contributing to OA progression. Additionally, transcriptomic shifts in immune cell subtypes were observed between low- and high-grade patients, providing new insights into OA severity. Finally, we showed that the metabolites highlighted by our cross-omics covariation analysis (without utilizing phenotype information) are enriched in markers of OA severity, validating our methodological strategy for addressing the challenge of high dimensionality. In summary, our work uncovers valuable mechanistic clues into individual variation in disease severity and also provides a methodological blueprint for large-scale multi-omics analysis of disease at cell type-resolution.

**Figure 1.**
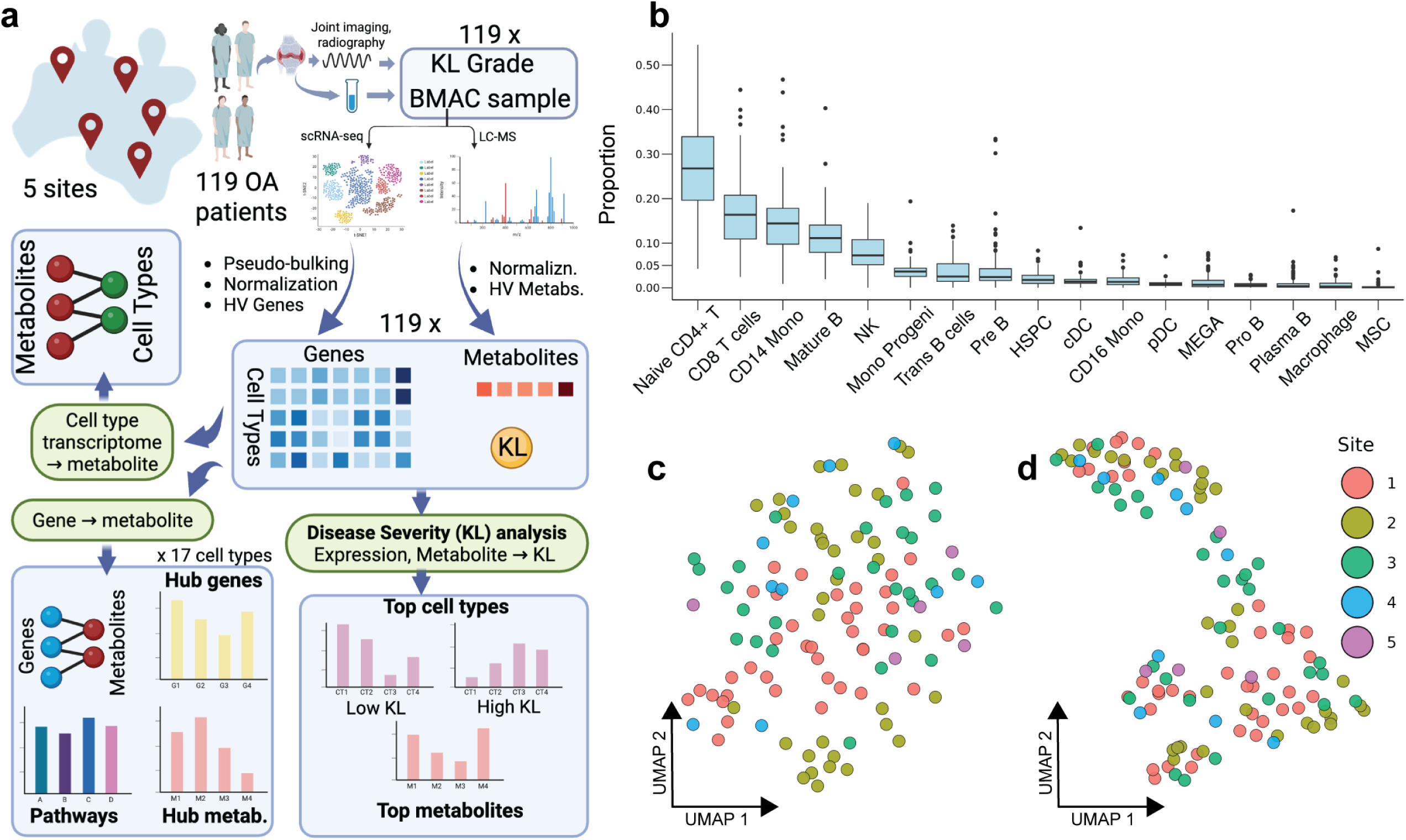
Data Description and Exploratory Data Analysis. **(a)** Schematic overview of dataset and analyses performed. BMAC samples and KL grades (OA severity) were obtained for 119 OA patients, and each sample was subjected to scRNA-seq and LC-MS profiling, yielding cell type-specific transcriptomic profiles and metabolomic profiles of each patient. These multi-omics data were analyzed to produce metabolite-gene associations for each cell type, highlighting key genes, metabolites and pathways, associating metabolites with cell types, and relating multi-omics profiles to KL scores. **(b)** Boxplots representing proportions of 17 cell types in BMAC samples across all patients. The cell types are sorted from left to right in decreasing order of median abundance. **(c, d)** UMAP plots of batch-corrected transcriptomics profiles (C) and batch-corrected metabolomics profiles (D). Each point represents a patient sample and is colored according to the site (hospital) where the sample was collected.

## RESULTS

### Metabolomic and cell type-resolved transcriptomic profiles of osteoarthritis patients

We generated LC-MS metabolomics profiles of bone marrow aspirate concentrate (BMAC) samples from 119 osteoarthritis (OA) patients enrolled in the MILES clinical trial for cell therapy [18]. These profiles were analyzed along with single-cell RNA-seq (scRNA-seq) profiles of the same 119 samples, from Chatterjee et al. [19]. The transcriptomics data revealed 17 cell types, and the average (pseudo-bulk, see Methods) transcriptome as well as relative abundance (proportion) of each cell type were calculated separately for each subject’s BMAC sample. We noted up to an order of magnitude difference among cell types in their relative abundances (**Figure 1B**), with lymphocytes such as naïve CD4 T cells, CD8 T cells, mature B cells and Natural Killer (NK) cells, as well as CD14 monocytes being the most frequent types. While the vast majority of cells found in BMAC are immune cells, the samples include, on average, ∼0.12% mesenchymal stem cells (MSC), ∼1.8% hematopoietic stem cells and progenitor cells (HSPC), and ∼0.67% megakaryocytes. Dimensionality reduction and visualization of the pseudo-bulk transcriptomes with Uniform Manifold Approximation and Projection (UMAP) shows no obvious separation among samples by sex, therapy response or site of sample collection (**Figure 1C**, **Supplementary Figures 1 A,B**). Similar UMAP visualization of (bulk) metabolomic profiles initially revealed a strong separation across sites, raising concerns about batch effects, but we remedied this using a batch effect correction procedure (**Figure 1D, Supplementary Figures 1 C,D**), after which there remained no clear segregation by sex, therapy response or site of sample collection.

### Covariation between metabolite and gene expression reveals key molecular factors in OA

Gene expression and metabolite abundance are known to influence each other [8]. We took advantage of the multi-omics compendium of BMAC samples to build a comprehensive map of gene-metabolite covariation across individuals, at cell type resolution. We focused on a set of 2000 highly variable genes and 1000 highly variable annotated metabolites (see Methods), discretized metabolite abundance using a percentile cutoff and tested each gene for differential expression between the samples defined by each metabolite’s presence or absence. (The test was repeated for each cell type.) This analysis identifies the extent to which a metabolite’s variation is associated with a gene’s expression variation in a cell type, ignoring the effects of varying cell type proportions.

The resulting map (**Supplementary Data S1A**) consists of ∼35k gene-metabolite pairs (∼1.7% of 2M pairs tested) on average in each cell type, with ∼20-30 genes found to have their expression in a cell type significantly associated (nominal p-value ≤ 0.005) with each metabolite (**Figure 2A**). (Also see **Supplement S1B** for results with adjusted p-values.) We noticed, for each cell type, that some genes and metabolites have significantly above-average number of associations. For instance, the map for macrophage cell type (**Figure 2B**) includes on average ∼38 gene associations per metabolite and ∼20 metabolite associations per gene, but the top 5% (50) metabolites have over 90 associations each and the top 5% (100) genes have over 60 associations each. We designated such genes and metabolites as “hubs” of the covariation map (**Methods**). Furthermore, we noted a dense interconnection between hub genes and metabolites, especially in the cell types Trans B cells, Pro B cells, CD14 monocytes (p-value < 0.01, **Methods, Supplementary Data S10**, **Supplementary Figure 2**), prompting us to delve into their functional interpretation. We also noticed a pronounced tendency for the hubs to be shared across cell types: the number of cell types that any gene/metabolite is a hub for has a significantly heavier tail than the randomly expected (Binomial) distribution; see **Figures 2C,D**. Based on these observations, we characterized hub genes/metabolites as “pan-cell type” if their average number of associations per cell type is large, and as “specific to a cell type” if they have many more associations in that cell type compared to other cell types on average (**Methods, Figures 2E,F, Supplementary Data S2**).

**Figure 2.**
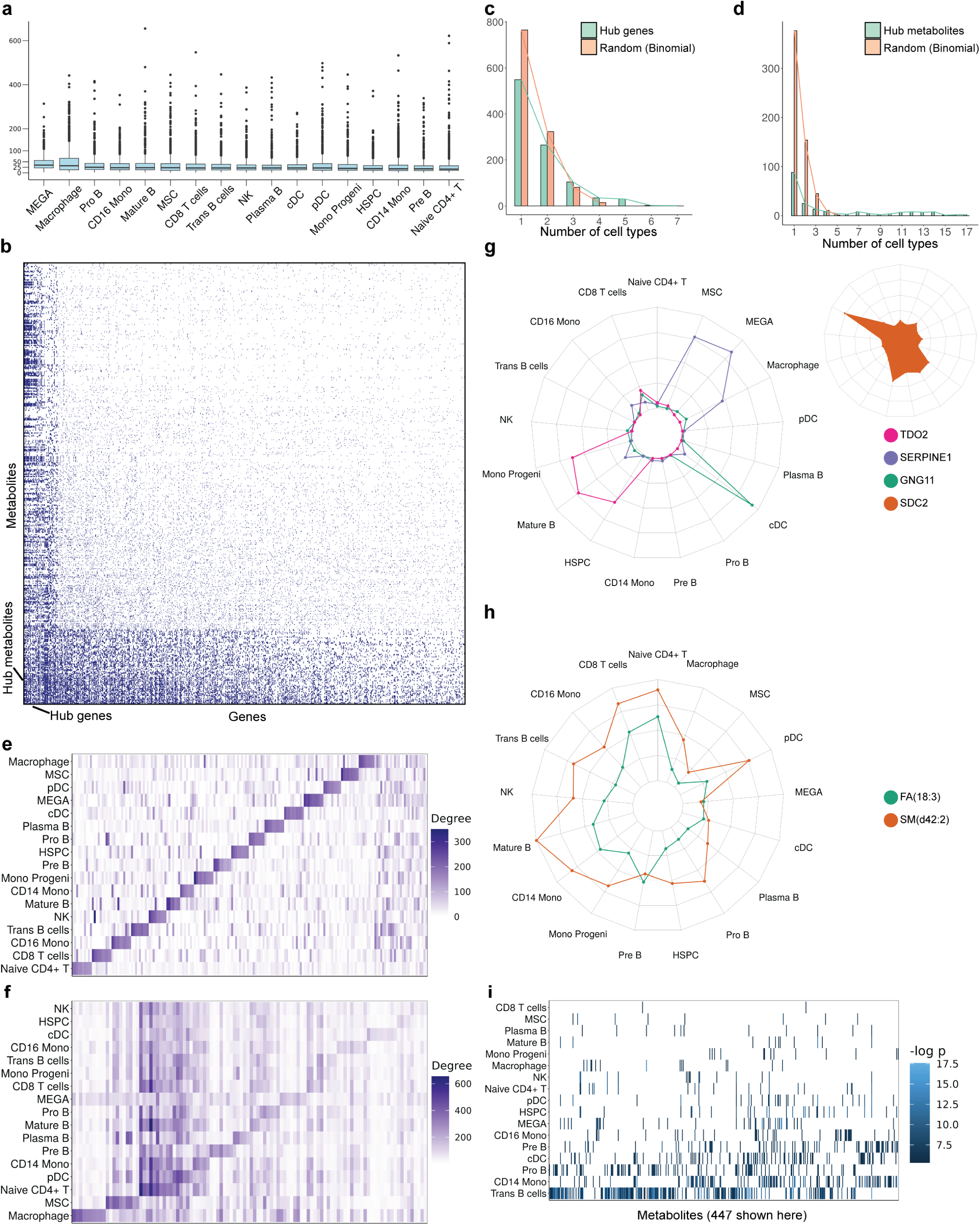
Covariation between metabolite and gene expression levels points to key molecular factors in osteoarthritis. **(a)** Boxplots showing the number of genes associated with each metabolite (at nominal p-value < 0.005) within each cell type. Each box shows median and 25^th^,75^th^ quartiles, along with outliers. We observe an average of 20-30 genes associated with each metabolite in every cell type. **(b)** Heatmap showing associations between 600 metabolites and 1200 genes in macrophage cell type. Each blue dot represents a significant association (nominal p-value < 0.005). There are ∼100 genes (leftmost columns) that are associated with at least 60 metabolites each, far greater than the average of ∼20 metabolite associations per gene. We call such genes hub genes (of macrophage in this case). Similarly, we observe that there are ∼50 metabolites (rows near the bottom) that are associated with at least 90 genes each, much larger than the average ∼40 gene associations per metabolite. We call such metabolites hub metabolites. (**c,d**) Green: Histogram of the number of cell-types for which a gene (c) or metabolite (d) is a hub. Orange: Distribution of Binomial random variable with size 17 and probability 0.05, which represents the null distribution corresponding to the histogram in green. (Only non-zero values have been shown.) **(e)** Heatmap showing the degree (number of metabolite associations) of each cell type-specific hub gene and of pan-cell-type hub genes in each cell type. **(f)** Heatmap showing the degree (number of gene associations) of each cell type-specific hub metabolite and pan-cell-type hub metabolite in each cell-types. **(g)** Spyder plots showing the degrees of cell-type specific hub genes (TDO2, SERPINE1, and GNG11) and pan-cell-type hub gene (SDC2) with strong literature evidence related to OA. **(h)** Spyder plot showing the degrees of cell-type specific hub metabolite (FA (18:3)) and pan-cell-type hub metabolite (SM (d42:2)). **(i)** Heatmap showing the strength of association (-log_10_(p-value)) between gene SDC2 and each of 447 metabolites across different cell types. SDC2 is an example of a pan-cell type hub gene and the plot shows that it is significantly associated with many metabolites in multiple cell types.

We next examined a selection of the hub genes in detail, focusing on the top 10 hubs specific to each cell type as well as the top 50 pan-cell type hubs (**Figures 2E,G**). For instance, the gene *TDO2* (Tryptophan 2,3-dioxygenase) is among hubs specific to mature B cells, with its expression in this cell type significantly associated with 186 metabolites, but with no associations in 11 (out of 16) other cell types. Interestingly, expression of TDO2, a key enzyme of the kynurenine pathway, in synovial fluid is a marker of osteoarthritis incidence and severity [20]. The kynurenine pathway has been shown to have a pro-inflammatory role in B cells [21] and is causally associated with osteoarthritis [22], and our finding about *TDO2* suggests a potential role for this gene in mature B cells in the context of OA. Another example is that of *SERPINE1* (Serine protease inhibitor E1), found to be a hub gene specific to megakaryocytes. There is evidence for both *SERPINE1* [23] and megakaryocytes [24] having potential roles in inflammatory joint diseases, so our finding raises the possibility that megakaryocyte-derived SERPINE1 plays a role in the inflammatory processes of OA. The gene *GNG11* (Guanine Nucleotide Binding Protein Gamma 11) was characterized as a hub gene specifically for conventional dendritic cells (cDC), its cDC expression levels being associated with 237 metabolites but fewer than 12 associations noted in 14 of the remaining 16 cell types. This gene has been suggested as a biomarker for OA-related inflammatory conditions [25] through functions in immune cell signaling, and cDCs are known to contribute to OA inflammation and immune response in the joint microenvironment [26], thus supporting our finding of *GNG11* as a cDC-specific hub gene for metabolic variation in OA.

In contrast to the above cell type-specific hubs, we also noted several genes as hubs across most cell types. One such pan-cell-type hub is the gene *SDC2*, significantly associated with over 20 metabolites in each of ten different cell types (**Figures 2G,I**). SDC2 (Syndecan-2), a transmembrane proteoglycan, regulates matrix metalloproteinases [27] whose dysregulation is a hallmark of OA [28], participates in cellular signaling pathways related to arthritis [29], has osteoblastic functions that affect bone remodeling [30] and is overexpressed in synovial membrane of OA patients [31]. Other notable pan-cell type hub genes associated with OA include *BMP3*, *ALPL*, *TPSB2*, *SULF1,* and *ALDH1A3* each of which is associated with at least 20 metabolites in at least eight cell types each (**Supplementary Figures 3 A-E** respectively). BMP3 plays a crucial regulatory role in bone mass maintenance [32], while ALPL (Alkaline Phosphatase) has been identified as a key player in bone development and remodeling [33, 34].

Analogous to hub genes, our analysis also revealed many hub metabolites (**Methods**, **Supplementary Data S3**), including cell type-specific hubs and pan-cell type hubs (**Figures 2F,H**). An example of a pan-cell type hub is the metabolite “SM(d42:2)” (**Supplementary Figure 3F**, Figure 2H), a sphingomyelin species that is known to be one of the most elevated sphingolipids in synovial fluid of OA patients [35]. We observed its abundance to be significantly associated with the expression of its synthesizing enzyme SGMS2 in CD14+ monocytes (p-value 0.0005) and macrophages (p-value 0.0016). This enzyme is central to homeostasis of ceramide [36] which is associated with pro-inflammatory signaling in rheumatic diseases [36] and in OA specifically [37]. Some metabolites were noted as being cell type-specific hubs, i.e., associated with many genes’ expression in one or few cell types. For instance, the polyunsaturated fatty acid FA(18:3) is associated with 42 genes in CD8+ T cells and 53 genes in Naïve CD4+ T cells (based on adjusted p-values at 0.05 significance level, **Supplementary Figure 3G, Supplementary Data S1B, Figure 2H)**. FA(18:3), also known as *α*-linoleic acid (ALA), and its metabolites have known immunomodulatory effects [38], can impact T cell activity [39], and are associated with reduced synovitis in OA knees [40].

In summary, we see extensive evidence of covariation between metabolite abundance and gene expression at cell type resolution, and many of these statistical associations point to genes and metabolites with known or expected roles in osteoarthritis. This analysis also catalogs a large number of less-studied but potentially functional factors of OA as hubs of the covariation network, that may serve as starting points for future studies.

### Gene set analysis identifies OA-related processes in distinct cell types

We subjected the hub genes of each cell type to gene set analysis using the Gene Ontology (GO) compendium, to extract systems-level insights from the gene-metabolite covariation map. We noticed that the hub genes of CD8 T cells are significantly enriched (p-value < 0.0005) in several OA-relevant biological processes (**Figure 3A, Supplementary Data S4)**, the strongest association being for the process “regulation of endocrine process”. Endocrine hormones like estrogens and parathyroid hormone influence bone metabolism and immune responses, which are important to OA [41], and endocrine disorders can exacerbate musculoskeletal issues, including OA [42]. Moreover, regulation of endocrine processes can modulate the differentiation of CD8 T cells and such differentiation in knee joints is known to be related to OA stage and compartment [43]. Notable among associations involving other cell types is that between hub genes of Naïve CD4 T cells and the process “negative regulation of stress-activated MAPK cascade”. Indeed, MAPK signaling plays an important role in function and differentiation of CD4 T cells [44] as well as regulation of osteogenic and chondrogenic differentiation [45], and MAPK inhibitors have a protective effect in early-stage OA [46]. Another prominent association identified is that between “zymogen activation” and (hub genes of) Progenitor B cells, both of which have relevance to OA [47, 48] but have not been studied in combination.

**Figure 3.**
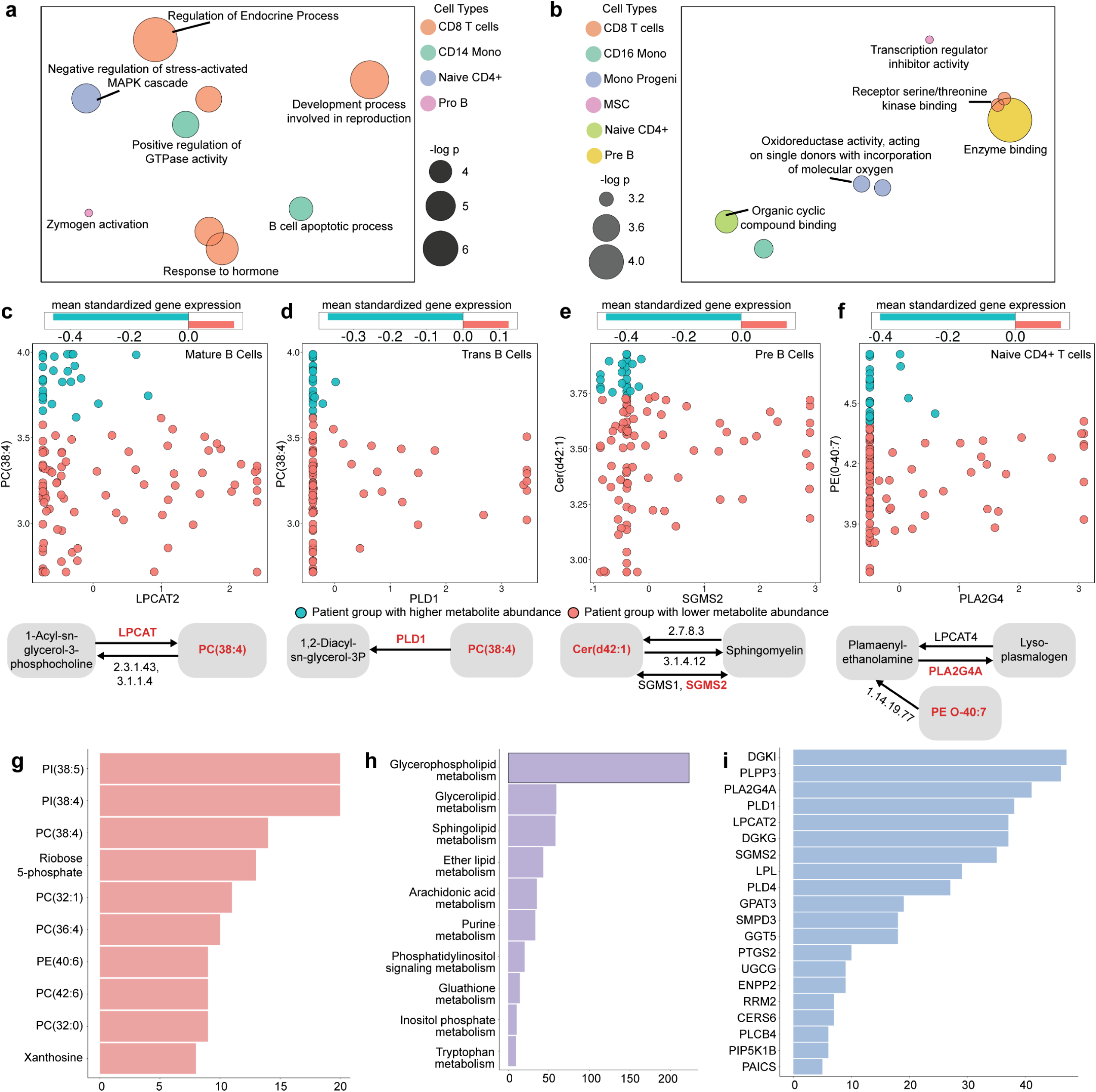
Gene set and pathway analysis of gene-metabolite associations in osteoarthritis. (a, **b)** REVIGO plots showing prominent Gene Ontology (GO) terms – biological processes (a) and molecular functions (b) – enriched in hub genes of different cell types. Colors represent cell types and circle size represents strength of enrichment (-log_10_(p-value)). Placement of circles has been calculated by the REVIGO tool that maps GO terms to a latent space based on similarity between GO terms. **(c, d, e, f)** Each panel depicts a gene-metabolite pair with significant statistical association in a particular cell type, and shows covariation of metabolite abundance with gene expression (scatterplot, middle) and a schematic diagram (bottom) of a reaction in KEGG pathway that includes the gene and the metabolite. In each panel, patient samples are shown divided into two groups (Methods) based on metabolite abundance (blue = higher metabolite abundance, red = lower metabolite abundance). Above each scatterplot is a bar plot that shows the mean standardized gene expressions in each group of patients, depicting a significant inter-group difference in mean gene expression. Pairs shown are: (c) metabolite PC(38:4) and gene LPCAT2 (in Mature B cells) based gene expression of LPCAT2, (d) metabolite PC(38:4) and gene PLD1 (in Trans B cells), (e) metabolite Cer(d42:1) and gene SGMS2 (in Pre B cells), (f) metabolite ‘PE(O-40:7)’ and gene PLA2G4 (in Naïve CD4+ T cells). **(g)** Bar plot showing the number of times each metabolite appears in the same pathway as an associated gene, aggregated across cell types. Only top 10 metabolites have been shown. **(h)** Bar plot showing the number of times each metabolic pathway harbors a significant gene-metabolite pair. Only top 10 pathways have been shown. **(i)** Bar plot showing the number of times each gene appears in the same pathway as an associated metabolite, aggregated across cell types. Only top 20 genes have been depicted.

Gene set analysis also revealed “molecular function” GO terms associated with hub genes of each cell type (**Figure 3B**). One of the strongest such associations was observed for “oxidoreductase activity, acting on single donors with incorporation of molecular oxygen”, enriched (p-value <0.0007) in hub genes of Monocyte Progenitor cells. Prior research suggests that oxidoreductase activity is associated with OA through reactive oxygen species (ROS) [49], which can promote macrophage polarization in OA [50] and influence cartilage degradation and joint inflammation [51]. Hub genes of CD8 T cells were enriched for “receptor serine/threonine kinase binding”, consistent with the widely recorded role of signaling pathways such as mTOR in T cells in OA onset and progression [52]. A direct assessment of biological pathways (**Supplementary Data S4**) revealed the TGF-beta signaling pathway as enriched in hub genes of CD8 T cells. This pathway plays crucial roles in CD8 T cell differentiation and function [53] and is important in the onset and progression of OA disease [54, 55].

Thus, systems-level analysis of hub genes highlights diverse and cell type-specific biological functions implicated in osteoarthritis pathology, including endocrine regulation, stress response modulation, and key signaling pathways, suggesting potential mechanisms and pathways for targeted therapeutic intervention.

### Incorporating pathway information yields a high-confidence map of metabolite-gene covariation

The gene-metabolite map includes many examples where the covarying pair belongs to the same metabolic pathway, e.g., the gene encodes an enzyme catalyzing a reaction involving the metabolite, thus offering a direct explanation for their covariation and increasing our confidence in their functional relationship. This “high-confidence map”, comprising on average 45 gene-metabolite pairs per cell type, is reported in **Supplementary Data S5**. For instance, the phopshatidylcholine PC(38:4) is significantly associated (p-value 1.7E-05) with Mature B cell expression of *LPCAT2* (lysophosphatidylcholine acyltransferase 2) (**Figure 3C**), which catalyzes the reacylation step of the Lands cycle, converting lysophosphatidylcholine (LPC) back to phosphatidylcholine (PC) [56], and facilitates membrane homeostasis of inflammatory cells [57]. The same metabolite, PC(38:4), is also associated with Trans B cell expression of PLD1 (Phospholipase D1) (**Figure 3D**), which catalyzes the hydrolysis of phosphatidylcholine (PC) to produce phosphatidic acid (PA) [58] and plays an important role in inflammation, bone demineralization and osteoclast differentiation in the context of rheumatoid arthritis [59, 60], suggesting similar but unexplored roles in OA. We found the ceramide cer(d42:1) to be strongly associated with Pre B cell expression of SGMS2 (Sphingomyelin Synthase 2) (**Figure 3E**), which is responsible for conversion of the ceramide to sphingomyelin [61]. Ceramide accumulation, as might result from dysregulation of this enzyme, plays a role in cartilage loss, a hallmark of OA [62]. These three examples illustrate associated gene-metabolite pairs where the gene and metabolite are known to participate in the same reaction. We also noted examples where the pair is separated by one reaction step in the same metabolic pathway. For instance, the phosphatidylethanolamine lipid species PE(0-40:7) was found to significantly covary with *PLA2G4* levels in Naïve CD4+ T cells (**Figure 3F**). PLA2G4 (cytosolic phospholipase A2 group IVA) catalyzes the hydrolysis of plasmenyl-ethanolamine, an ether-linked phosphatidylethanolamine that is produced from PE(0-40:7) [63], and is associated with OA progression [64]. In all four of these examples (Figures 3C-F), higher enzyme expression was associated with lower metabolite abundance. Nearly 72% pairs in the high-confidence map follow this trend.

In many cases, a metabolite was found to covary with more than one gene within its metabolic pathway, allowing us to rank metabolites by the frequency of such high-confidence relationships. For instance, the four examples above included two genes (*LPCAT2*, *PLD1*) that covary with PC(38:4); in fact, the high-confidence map includes 14 associations (spanning multiple cell types) involving this metabolite. Overall, phosphatidylinositols (PIs) and phopshatidylcholines (PCs) were among the most frequent metabolites in the high-confidence map (**Figure 3G)**. Phosphatidylinositols play a significant role in OA pathogenesis and progression, primarily through the phosphatidylinositol 3-kinase (PI3K)/Akt signaling pathway, which is involved in cartilage degradation, subchondral bone dysfunction and synovial inflammation [65]. Phopshatidylcholine metabolism has been linked to knee cartilage volume loss in OA [66]. Another metabolite highlighted by the analysis, Ribose 5-phosphate, is a key product of the pentose phosphate pathway (PPP), which is important for chondrocyte survival [67] and is significantly up-regulated in OA [68].

The high-confidence map included over 200 pairs from “glycerophospholipid metabolism” pathway, by far the highest among all metabolic pathways (**Figure 3H**). Metabolomic studies have reported glycerophospholipid metabolism as strongly associated with OA [69] and one study found the spatial distribution of glycerophospholipids to be correlated with hypertrophic, inflamed or vascularized areas in OA synovium [70]. Glycerolipid and Sphingolipid metabolism were the second and third most represented pathways in the high-confidence map, both of which have some evidence of roles in OA [69].

Turning our attention to genes ranked by their frequency in the high-confidence map (**Figure 3I**), we noted *DGKI* (Diacylglycerol Kinase Iota) and *PLPP3* (Phospholipid Phosphatase 3) to be the most represented, with over 45 associations each. Both are enzymes involved in lipid signaling pathways [71, 72], which play critical roles in cartilage-related diseases [73]. Other highly ranked genes include *PLA2G4*, *PLD1* and *LPCAT2*, which are known to be associated with OA [64] or with processes important to OA [57, 59, 60].

To summarize, focusing on covarying metabolite-gene pairs within the same metabolic pathway highlights mechanistically grounded relationships and identifies key molecular players in OA pathogenesis, offering valuable insights into disease mechanisms and potential therapeutic targets.

### Cell types associated with metabolic variation in OA

The single-cell resolution of transcriptomics data presented us with the unique opportunity to determine which cell types, if any, are most strongly associated with inter-individual variation of a metabolite in the context of OA. Variations in a cell type’s molecular contributions to a tissue arise in part from changes in its molecular contents and in part from changes in the cell type’s proportion in the tissue. We examined each of these sources of variation separately vis-à-vis their relationship to metabolomic profiles of individuals in the OA cohort.

In our first analysis, we asked how well a cell type’s global transcriptome predicts a metabolite’s abundance, using a Machine Learning (Random Forest) model (**Methods**). We observed (**Figure 4A**) that 70 metabolites can be linked to a specific cell type’s transcriptome (balanced prediction accuracy ≥ 0.65 when using that cell type’s transcriptome). For example, the abundance of “L-Glutathione(reduced)” can be predicted using the transcriptome of Plasma B cells with balanced accuracy 0.65 (Binomial test p-value 0.03), but far less accurately using other cell types (**Figure 4B**), thus suggesting a link between this metabolite and cell type pair. This predicted link is supported by previous findings regarding roles of L-Glutathione in B cell development, function, and survival [74] as well as by its recorded protective role in OA via neutralization of ROS [75] and other mechanisms [76, 77]. Another illustrative case of metabolite-cell type association is that of “TG(49:1)” (a Tri-glyceride compound) that is predictable specifically using the transcriptome of NK cells (balanced accuracy 0.68, p-value 0.01, **Figure 4C**). Supporting this finding, triglycerides have been shown to suppress NK cell function via metabolic reprogramming [78] and are strongly correlated with OA risk [79]. Several such examples of metabolite-cell type associations were observed for most other cell types (**Supplementary Data S6**). On the other hand, we also observed 22 examples of metabolites that can be linked to multiple (four or more) cell types (**Figure 4A**). An example is “Metformin”, predictable (balanced accuracy >= 0.65) using transcriptomes of 16 cell types (**Figure 4D**). Metformin has been shown in preclinical studies to inhibit pro-inflammatory cytokines and matrix metalloproteinases while promoting autophagy in chondrocytes [80] [81]. Another example of a pan-cell type association is that of the metabolite PE(O-38:5), which predictable by transcriptomes of 11 cell types (**Figure 4E**). This ether-linked phosphatidylethanolamine has been previously reported as one of the most abundant lipids in connective tissue compared to adipose tissue, in a study of OA and cartilage defect patients [82].

**Figure 4.**
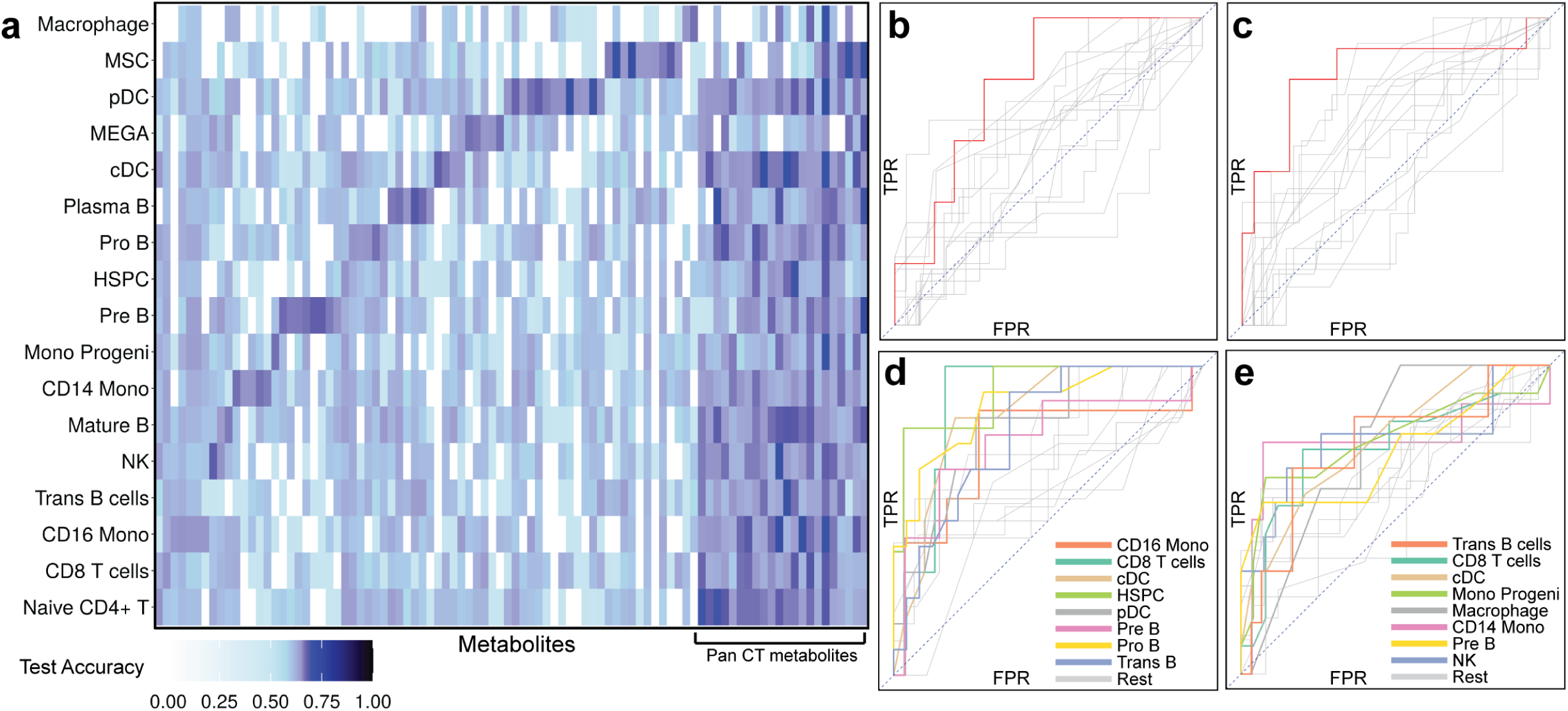
Cell types associated with metabolic variation in OA. **(a)** Heatmap showing the transcriptome-based predictability (balanced test accuracy, random expectation = 0.5) of metabolite abundance, for select metabolites (columns) in each cell type (row). The accuracy is that of a machine learning model (random forest) trained to predict metabolite abundance using transcriptome of each cell type. Columns near the right (“Pan CT metabolites) represent metabolites with high predictability in multiple cell types, while remaining columns represent metabolites with high predictability in specific cell types. **(b, c)** Receiver Operating Characteristic (ROC) curves for prediction of two select metabolites based on transcriptome profile of each cell-type, where each line represents a cell type. X-axis and Y-axis show False Positive Rate (FPR) and True Positive Rate (TPR) respectively. In each panel, one curve (shown in red) represents the cell type for which the test accuracy is significantly larger than that of other cell types (curves shown in gray), illustrating that the transcriptome-based predictability of the metabolite’s abundance is cell type-specific. Figure b shows results for metabolite “L-Glutathione(reduced)” and highlighted ROC curve is for Plasma B cells, while figure c shows results for metabolite “TG(49:1)” and highlighted ROC curve is for NK cells. **(d, e)** ROC curves for prediction of two select metabolites based on transcriptome profiles of each cell type. X-axis and Y-axis show False Positive Rate (FPR) and True Positive Rate (TPR) respectively. The colored ROC curves correspond to the eight cell types where the test balanced accuracies are larger than 0.65 while the curves for remaining cell types are shown in gray, illustrating that these metabolites are “pan cell type” predictable. Figure d shows results for metabolite “Metformin” while figure e corresponds to “PE(O-38:5)”.

We also tested for metabolite-cell type associations based on the statistical association between the metabolite’s abundance and the cell type’s abundance (proportion relative to other cell types). Less than 9% of the metabolites can be predicted more accurately using cell type’s proportion than using any one cell type (**Supplementary Data S6**), suggesting that metabolic variation arises predominantly from varying transcriptomic profile of a cell type rather than its abundance in the tissue.

### Cell type-specific multi-omics footprints of OA severity

The analyses thus far examined relationships between metabolomics and transcriptomics profiles of OA patient samples, but did not explicitly model any phenotypic variables. We now asked how either of these omics profiles varies in predictable ways with OA severity, recorded in the form of Kellgren/Lawrence (KL) grades, which in this cohort take the values of 2 (low), 3 (medium) and 4 (high). To ensure adequate sample sizes, we compared the KL-2 (low) group with others, and separately the KL-4 (high) group with others.

First, we trained ML models (Random Forest) to distinguish individuals in a severity group using their cell type-specific transcriptomic profiles (**Methods**). We noted that these models performed more accurately (on unseen test data) when trained using only hub genes (of a cell type) obtained from the covariation analysis, compared to using all genes (**Supplementary Figure 4**). This provides concrete evidence in support of our methodological premise – that gene-metabolite covariation analysis prioritizes genes that are relevant to the phenotype, even though phenotypic variables were not explicitly modeled in that analysis (due to high dimensionality).

The KL-4 (high grade) group was most accurately distinguished (**Figure 5A**) using expression data from CD14 monocytes (AUROC 0.72, compared to random baseline of 0.50) and Monocyte Progenitor cells (AUROC 0.71), significantly better than with any other cell type (**Supplementary Data S7**). On the other hand, the KL-2 (low grade) group was best classified (**Figure 5B**) by the transcriptomes of CD16 monocytes (AUROC 0.79), MSCs (AUROC 0.74) and Plasma B cells (AUROC 0.73), significantly better than other cell types (**Supplementary Data S7**). Notably, neither group is distinguishable using bulk transcriptomic data (AUROC of 0.56 and 0.55 respectively), underscoring the cell type-specificity of transcriptomic changes associated with OA severity.

**Figure 5.**
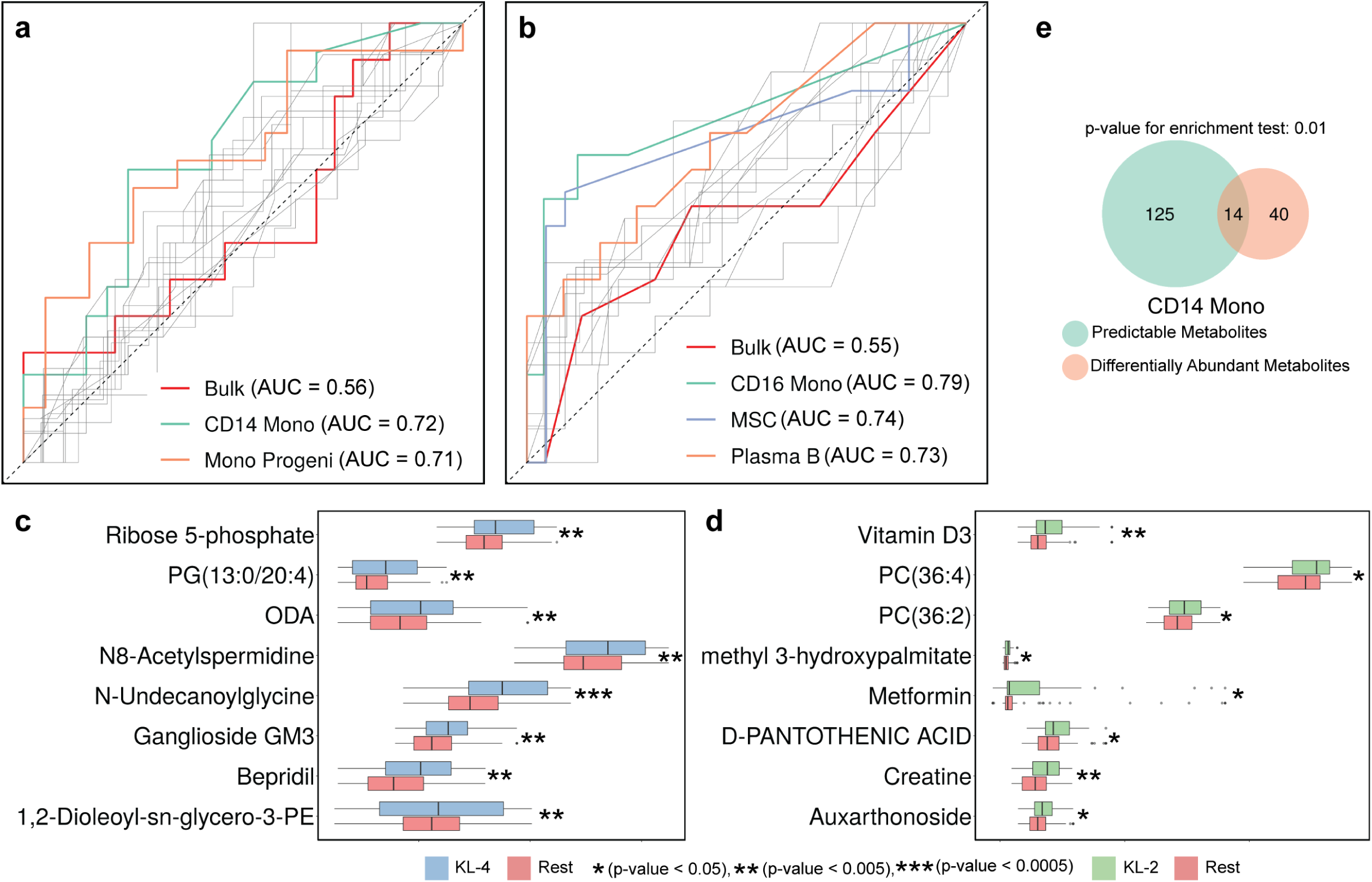
Cell type-specific multi-omics footprints of OA severity. **(a, b)** ROC curves for random forest-based prediction of KL grades using each cell type’s transcriptome (KL-4 vs KL- 2,3 in figure a and KL-2 vs KL-3,4 in figure b). The three ROC curves (cell types) with the highest area under curve (AUC) are highlighted, revealing the cell types with strongest global transcriptomic footprints of OA severity. The ROC curve of a predictor that uses bulk transcriptome is shown in red, while all other ROC curves are shown in gray. **(c, d)** Boxplots of metabolite abundance in KL-X group vs the rest (KL-4 on the left and KL-2 on the right). 8 metabolites with the strongest p-values are shown. **(e)** Venn diagram showing the intersection between the differentially abundant metabolites (KL-4 vs the rest, with more abundance in KL-4 group) and the metabolites that are predictable from pseudo-bulk transcriptome of CD14 monocytes. There is a significant enrichment (at 0.05 significance level) based on hypergeometric test.

Cell types with the strongest predictive power offer an interesting glimpse into disease progression: we noted a marked shift from CD16 monocytes to CD14 monocytes as the transcriptomic predictor of low and high grade respectively. This mirrors previous findings that CD14^+^CD16^−^ monocytes are the main precursors of osteoclasts in rheumatoid arthritis [83], that their proportion in synovial fluid is predictive of the Knee Injury and Osteoarthritis Outcome Score (KOOS) [84], and that CD14 monocytes play an important role in OA aggravation through inflammatory cytokine secretion [85]. The predictive power of mesenchymal stem cell (MSC) transcriptomes for the low grade group (AUROC = 0.74) but not the high grade group (AUROC = 0.55, close to the random baseline) is consistent with the demonstrated role of MSCs in pain and symptom reduction in OA clinical trials [86].

Next, we probed the metabolomic signatures of each severity group and identified 54 (resp. 17) metabolites significantly associated with the KL-4 (resp. KL-2) group using a test of different proportions (nominal p-value < 0.05, **Methods, Figures 5C,D Supplementary Data S8**). We asked if these phenotype-related metabolites could have been identified by the cross-omics analysis presented in earlier sections. To answer this, we identified (Supplementary Data S6) the metabolites whose abundance in patient samples can be accurately predicted using the pseudo-bulk transcriptome of CD14 monocytes, the cell type most relevant to the KL-4 group (see above). Fourteen of the 54 KL-4-associated metabolites were included in the prioritized list of 139 metabolites from the cross-omics analysis, a significant enrichment (Hypergeometric test p-value 0.01, **Figure 5E, Supplementary Data S9**). This result furnishes additional evidence that transcriptomics-metabolomics covariation analysis can reveal important molecular footprints and mechanisms of OA.

The above-mentioned short list of 14 metabolites that are identified both by cross-omics analysis and by differential abundance analysis for the KL-4 group provide exciting opportunities for targeted follow-up experiments in the future, since they are supported by two complementary types of analysis. Two of these metabolites are especially interesting: Cer(d40:0)M+CH3COOH, a ceramide, and SM(d42:2), a sphingomyelin lipid. Ceramide is a lipid messenger involved in apoptosis and matrix degradation in cartilage [62], both of which are closely related to OA. Sphingomyelin has been shown to be correlated with OA pathology and pain levels in OA [87], and the particular 42 carbon species SM(d18:1_24:1) is reported to have significantly elevated levels in OA [70].

## DISCUSSION

Our study uses a multi-omics approach to investigate molecular mechanisms underlying osteoarthritis (OA). We analyzed data on bone marrow aspirate concentrate samples from 119 OA patients participating in a clinical trial, constructing detailed maps of gene-metabolite relationships in each of 17 distinct cell types. This analysis revealed cell type-specific as well as pan-cell type hub genes and metabolites, highlighting their roles in inflammatory processes, cartilage homeostasis, and metabolic dysregulation. Focusing specifically on gene-metabolite pairs that belong to the same pathway helped us identify critical pathways such as phosphatidylinositol, phosphatidylcholine, sphingolipid, and glycerophospholipid metabolism, along with lipid signaling, to be central to OA. Going beyond univariate analysis involving individual gene-metabolite pairs, we used Machine Learning models to relate the entire transcriptome of a cell type to each metabolite, thus identifying specific cell types most strongly associated with a metabolite’s inter-individual variation. We supplemented our cross-omics analysis with data on disease severity in the subjects, observing a shift in immune cell subtypes between low- and high-grade OA patients. Finally, we showed that focusing on cross-omics statistical signals (genes-metabolite and whole transcriptome-metabolite relationships) is an effective strategy for dealing with the challenges faced in predicting phenotypic variation and its biomarkers from high dimensional multi-omics data.

A key strength of our approach is the high-resolution mapping of cell type-specific molecular networks. The use of single-cell RNA sequencing and identification of 17 distinct cell populations allowed us to decouple the contributions of each cell type to the overall metabolic landscape of OA. The importance of metabolic characterization in a cell type-specific manner has been argued by other studies [88], but progress has been limited. Our systematic examination of the reconstructed molecular networks revealed cell type-specific omics signatures, such as the mature B cell-specific hub gene TDO2 and the CD8/CD4 T cell-specific hub metabolite FA(18:3), as well as more broadly acting (pan-cell type) hubs such as SDC2 and SM(d42:2). In several cases the cell type-specific roles of identified genes have been reported in the literature, e.g., B cell-related pro-inflammatory functions of the kynurenine pathway (that includes TDO2) or the immunomodulatory effects of FA(18:3) on T cell activity in OA, but many known/hypothesized molecular associations of the disease do not have cell type assignments and our findings furnish a large compendium of candidates to be tested in specific cellular contexts.

The combination of single-cell transcriptomics with metabolomics in a clinical trial cohort is a novel aspect of our study. There is some precedence of gene-metabolite covariation analysis at population level in the context of cancer, where data from “The Cancer Genome Atlas” were analyzed to reveal enzyme-substrate interactions and hub metabolites linked to immune cells [13]. Similarly, Li et al. [12] profiled metabolomes of ∼900 cancer cell lines and identified associations between metabolites and genetic/epigenetic variables. Our work applies a similar investigative paradigm to OA, while also advancing it by introducing cell type resolution to the data and analysis. Prior OA studies often focused on genetic, transcriptomic, or epigenomic markers [89], and also reconstructed co-expression networks and modules [90], but comprehensive gene-metabolite analysis at cell type level has been missing.

Our “high-confidence map” of gene-metabolite pairs active within the same metabolic pathway is inspired by the “proximal gene-metabolite interactions” identified by Benedetti et al. [13] in a cancer context, and shortlists many predictions of enzymes as major determinants of specific metabolite pool sizes, in specific cell types. For instance, the map highlights the interplay between phosphatidylcholine metabolism and key enzymes such as LPCAT2 and PLD1, corroborating the established involvement of glycerophospholipid metabolism in cartilage degradation and joint inflammation [4, 91].

Our cell type-specific severity-prediction models illuminate some of the dynamic changes associated with OA progression. The distinct transcriptomic signatures of CD14+ versus CD16+ monocytes and the differential predictive power of mesenchymal stem cell (MSC) profiles in low-grade versus high-grade OA underscore the nuanced roles that individual cell types play in disease progression. These findings align with earlier reports implicating monocyte subsets in joint inflammation and osteoclastogenesis [83, 85], while also emphasizing the utility of cell type-resolved analyses for capturing clinically relevant heterogeneity. At the same time, our main goal was not to build a predictive or diagnostic model for OA progression. This is a daunting challenge even with > 100 sample size analyzed here, due to the very high dimensionality of omics profiles and due to a high level of intra-class variability both at the molecular and phenotypic level [19]. Thus, our emphasis was on molecular relationships underlying variations in the cohort, with the caveat that these associations are not explicitly assigned importance scores for phenotype prediction. Xie et al. [10] note that cataloging gene-metabolite covariation has another practical advantage – the ability to impute metabolite levels in clinical samples, which they achieved using a novel Bayesian framework.

Our statistical analyses explore relationships at different levels of biological organization, including associations between one gene and one metabolite (univariate analysis) as well as models relating an entire transcriptome (of a cell type) to individual metabolites (multi-variable analysis). Univariate analysis was supported by a moderate sample size (∼120 patient samples) but suffers from a multiple hypothesis testing burden, which we addressed by focusing on hubs (rather than individual gene-metabolite pairs) and by filtering gene-metabolite pairs for co-membership in a characterized metabolic pathway. Multivariable analysis (e.g., when predicting a metabolite’s abundance or patient phenotype from entire transcriptomes) poses the challenge of high dimensionality, and we addressed this via feature selection strategies or, in the case of phenotype prediction, by using hub genes as selected features. Our choice of Random Forest (non-linear) models was based on preliminary assessments comparing these to linear or logistic models. All analyses were performed after discretization of metabolite abundance levels into high (75^th^ percentile and above) versus low levels. Our motivation here was mainly to reduce the impact of technical noise in metabolite abundance measurements, providing more robust correlations with gene expression levels, especially during multi-variable analysis where classification (Random Forest) was performed instead of regression. Our multi-omics analysis separately examined the effects of average expression level in a cell type and that of cell type proportions. We preferred this to examining their joint effect (as reflected in a product), in order to deconvolve the contributions of these two complementary sources of transcriptomic variation, with concomitant ease of interpretation.

The cross-sectional nature of the omics data at the heart of our study, and their origin in a single clinical trial cohort suggest some caution regarding the generalizability of our findings. Future studies employing longitudinal sampling and independent cohorts will be important for validating the reported associations, along with functional validation of molecular mechanisms through appropriate perturbation experiments.

## METHODS

### Data Generation

#### Metabolome

Broad spectrum LC-MS was performed on cell pellets of 1 million nucleated cells per patient sample, using the Thermo Fisher Scientific Vanquish Horizon UHPLC coupled with Orbitrap ID-X Tribrid Mass Spectrometer system. In order to detect a broad range of metabolites, samples were analyzed using Reverse Phase (RP) chromatography as well as Hydrophilic Interaction Liquid Chromatography (HILIC), in both negative and positive mode. A pooled QC sample was created by taking an aliquot from each patient sample and combining these, and the pooled QC sample was injected at least every 10 injections throughout the run and used to correct for instrument drift. A blank sample was created through the same process as the cell samples but excluding cells and was analyzed at the beginning of the batch. All samples were run in the same batch to avoid batch effects from the data acquisition step, and the run order for the samples in the batch was randomized. Data processing and basic analysis was performed using Compound Discoverer V3.1 software (ThermoFisher Scientific).

### Data Preprocessing

#### Transcriptome

We utilized single cell sequencing data generated by Chatterjee et al. [19], recording expressions of 27930 genes in 119 BMAC patients in total from 5 different sites. Seurat tool was used to remove any potential batch effects and to perform cell type annotations for each cell. The Seurat tool chooses a smaller set of highly variable genes to accomplish the aforementioned tasks and in this case a set of 2000 highly variable genes was used. A highly variable set contains the genes that have the largest variance across the cohort of patients. The same set of 2000 genes has been used for all analyses in this paper. The cells were categorized into 17 different cell types. The above preprocessing steps are as reported in [19].

The average expression of each gene from all the cells of a patient was recorded to construct a bulk expression vector for the patient. Similarly, the average expression of each gene from all the cells of a cell type was recorded to construct cell type-specific pseudo-bulk expression vector. Therefore, each patient has a bulk expression vector and 17 cell type specific pseudo bulk expression vectors.

To reduce the impact of outliers, the expression values of each gene (bulk as well as pseudo-bulk) were saturated at the 95^th^ quantile of that gene. In other words, for a given gene, the expression values greater than or equal to the 95^th^ quantile were changed to the 95^th^ quantile value. Furthermore, the expression of each gene (bulk as well as pseudo-bulk) was centered to mean zero and scaled to have standard deviation 1.

### Metabolome

Broad spectrum untargeted LC-MS analysis was used to record bulk abundance of around 7000 polar and non-polar metabolites in each patient sample, as described above. Data was processed using Compound Discoverer v3.1 to remove background and correct for instrument drift. Following this, values for each metabolite peak were quantified as the normalized area under the curve (AUC). Annotation of the dataset was performed by MS2 spectral matching to a local spectral database as well as online sources. Mass, retention time, and isotopic pattern were matched to database entries. This resulted in ∼1400 of the detected metabolites being annotated with names. To remove any potential batch effect, Seurat tool was utilized to select 1000 most variable metabolites among these named metabolites. The same set of 1000 (named) metabolites has been used for all analyses in this paper.

To reduce the impact of outliers, the abundance values of each metabolite was saturated at the 95^th^ quantile of that metabolite, as was done for genes. Furthermore, the expression of each metabolite was centered to mean zero and scaled to have standard deviation 1.

### Metabolite-Gene Pair Associations

The aim is to find associations between metabolites and genes at a cell type level. For a given metabolite, the patient samples were divided into two groups based on metabolite abundance. The samples with metabolite abundance greater than or equal to the 75^th^ quantile were assigned Group 1 and rest of the samples were assigned Group 2. Next, for a given cell type, the pseudo-bulk gene expressions of each gene were used to perform differential expression (DE) analysis between the two groups of patients. For each gene, the null hypothesis states that the expression of the gene is same in the two groups whereas the alternate hypothesis states that the gene expressions between the two groups are different. A two-sample t-test was utilized to accomplish this task. Consequently, a p-value for the strength of association was obtained for each metabolite-gene pair in a cell type.

### Justification for discretization of metabolites

The exact metabolite abundance values as measured by peak area are relatively noisy and the relative order of metabolite abundances between samples is a more reliable measure for the purposes of comparison. Therefore, we divided patients into two groups: one with higher metabolite abundance (above the 75th percentile) and the other with lower abundance (below the 75th percentile). This threshold was chosen to ensure a sufficient number of samples in both groups for comparative analysis.

Alternatively, a rank-based correlation analysis between a gene and metabolite pair could be performed. However, since many patients have either very low gene expression or very low metabolite abundance, assigning equal weight to all samples could be problematic because such samples can easily dominate the correlation results. Grouping patients into high vs. low abundance categories provides a more intuitive and interpretable approach for the comparative analysis.

### Hub genes and Hub metabolites

In each cell type, on average, we observed 50 metabolites to have over 70 gene associations each. In contrast, other metabolites have, on average, 25 gene associations each. Therefore, we designated top 50 metabolites (in terms of the number of genes associated with them) as hub metabolites.

In each cell type, on average, we observed 100 genes to have over 60 metabolites associated each. In contrast, other genes have, on average, 15 metabolite associations each. Therefore, we designated top 100 genes (in terms of the number of metabolites associated with them) as hub genes.

#### Cell-type-specific hubs

We observed certain hub genes and metabolites to be specific to each cell type. To make the definition of a cell type-specific hub more concrete, we defined a score for specificity of a hub gene or metabolite to a cell type *i* as follows: 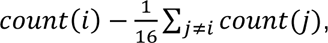 where *count*(*k*) is the number of associations of the gene/metabolite in that cell type. In other words, this is the difference between the number of associations of the gene/metabolite in a cell type and the average number of associations in other cell types. Then we sort and select the 10 genes/metabolites with the highest values of this score. This set serves as cell type-specific hub genes or metabolites.

#### Pan-cell-type hubs

We observed several hub genes or metabolites to have large numbers of associations (characteristic of a hub) in multiple cell types. To objectively define such “pan-cell-type hubs”, we performed the following steps. For each cell-type, we ranked genes/metabolites based on the number of associations (lower rank means more associations). In the next step, we combined the ranks of each gene/metabolite across all the cell types by calculating geometric means of their ranks. Based on the calculated geometric mean, we ranked the genes/metabolites again to obtain a combined rank (lower combined rank corresponds to overall lower rank across cell-types). We selected the top 50 genes/metabolites according to the combined rank as pan-cell-type hub genes/metabolites.

#### Cluster Scores for interconnection between hub genes and hub metabolites

We used the bipartite function from NetworkX library in Python to calculate the density score between two sets of nodes – hub genes and hub metabolites. Furthermore, we randomly pick 100 non-hub genes and 50 non-hub metabolites to calculate the density score, and repeat this step 200 times. We define the p-value as the fraction of times the density score of non-hub interconnections are larger than the density score of hub interconnections (**Supplementary Data S11)**.

#### REVIGO Plots

We used the REVIGO tool [92] to plot the results obtained from gene set enrichment analysis from Gene Ontology compendium. The tool represents the GO terms in two-dimensional space so that terms that are similar appear closer in the plot. The size of the circle represents the strength of enrichment between the given set of hub genes and the genes from the GO term.

### High-confidence map description

The map was constructed for each cell type separately, as follows. For each metabolite, we first identified the set of associated genes based on cell-type-specific pseudo-bulk gene expression profiles at a nominal p-value threshold of 0.005 (see above). We then checked which of these genes shared a metabolic pathway with the metabolite. When such a match was found, we recorded a high-confidence quadruplet – (metabolite, gene, cell type, pathway), indicating that the gene and metabolite participate in the same pathway in a cell-type-specific context. To obtain pathway information, we web-scraped the Metabolomics Workbench [93] to retrieve all pathways involving the metabolite and collected their corresponding KEGG IDs. Using the KEGG API [63], we accessed the list of genes involved in each pathway and intersected this list with the genes associated with the metabolite in that specific cell type. This intersection provided a high-confidence mapping between metabolites and genes. We repeated this process across all cell types.

### Metabolite-Cell Type Associations

The aim is to find a set of metabolites associated with each cell type, based on a transcriptomic profile of that cell type. For a given metabolite, the patient samples were divided in two groups based on metabolite abundance. The samples with the metabolite abundance greater than or equal to the 75^th^ quantile were assigned Class 1 and rest of the patients were assigned Class 2. For a given cell type, its pseudo-bulk transcriptome was used to predict these two classes via a Random Forest classifier.

To assess the performance of the Random Forest model, 70% samples were randomly assigned to the train set and the remaining 30% samples were assigned to the test set. Within the training set, a set of significant DE genes (0.05 significance level) was obtained via a two-sample t-test. The pseudo-bulk expression levels of these DE genes were used as the input features for the Random Forest classifier. The number of trees and the tree-depth of the classifier were chosen based on cross validation within the training set, implemented using RSCV function from sklearn. The best Random Forest classifier was then used to predict the class labels on the test set. The input features of the test set are same as the significant DE genes obtained during the training step. Subsequently, a balanced accuracy was obtained on the test set. We repeated the steps of splitting the data into train-test set, training the model and testing the model 100 times to obtain 100 balanced accuracies. Next, we created a 95% confidence interval based on these balanced accuracies centered around their mean 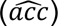 and standard deviation calculated based on the mean of the 100 balanced accuracies 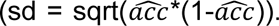 If the lower end of the accuracy interval is less than 0.5, we report the mean of the 100 accuracies. On the other hand, if the lower end of the accuracy interval is strictly greater than 0.5, we report the median of top 20 accuracies.

### Transcriptome-KL associations

We used the pseudo-bulk transcriptome for each cell type to train a Random Forest classifier to classify patient samples in the KL-X group vs those in the other two KL groups (X is 2 or 4 and represents KL score). For each cell-type, we only consider those genes as input features which we had designated as hub genes for that cell type based on covariation analysis.

To assess the performance of the Random Forest model, 70% of the patient samples were randomly assigned to the train set and the remaining 30% of patient samples were assigned to the test set. Within the training set, a set of significant DE genes (0.05 significance level) was obtained via a two-sample t-test. The pseudo-bulk expression of these DE genes was used as the input features for the classifier. The number of trees and the tree-depth of the random forest classifier were chosen based on cross validation within the training set, which was implemented using RSCV function from sklearn. The best random forest classifier was then used to predict the class labels on the test set. The input features of the test set are same as the significant DE genes obtained during the training step. We repeated the steps of splitting the data into train-test sets, training the model and testing the model 30 times and selected the run for which balanced accuracy is the median across 30 runs. We then plot the ROC curve (on test data) for this selected run of the classifier and report area under ROC curve (labelled as AUC in results).

### Metabolite-KL associations

To identify associations between each KL score (specifically KL 2 and KL 4) and individual metabolites, we carried out the following steps. First, for each metabolite, we created a binary vector by categorizing samples into two groups: Group 1 if the metabolite expression was above the 75th percentile, and Group 0 otherwise. This allowed us to represent metabolite abundance in binary terms. Next, we calculated two proportions to evaluate the distribution of high-abundance samples across KL score categories. For example, considering KL=2, we computed: (a) the proportion of Group 1 (high-abundance) individuals among KL=2 patients, and (b) the proportion of Group 1 individuals among all other patients. We then conducted a proportion test (prop.test) to compare these proportions. The null hypothesis posited that the proportion of patients with high metabolite abundance is the same in KL2 patients and the rest, while the alternative hypothesis suggested that the proportion is greater in KL2 patients.

### Overlap between Transcript-based-predictable metabolites and DA metabolites

For each cell type, we first selected metabolites that had test accuracies greater than 0.6. To ensure specificity, we excluded any metabolites for which the test accuracies from confounders were greater than or equal to 0.6. After this filtering step, we identified the final set of relevant metabolites by taking the intersection between these selected metabolites and the differentially expressed (DE) metabolites.

## Supporting information

Supplementary Data Description

## DATA AVAILABILITY

The bone marrow aspirate concentrate (BMAC) single-cell RNA sequencing dataset generated in this study has been deposited in the Gene Expression Omnibus (GEO) under accession number GSE274018 and is also available in the Sequence Read Archive (SRA) under project number PRJNA1144164. The raw metabolomics data are being prepared for upload to the NIH Metabolomics Workbench and will be made publicly available prior to publication of this manuscript. The full BMAC cohort dataset, including additional single-cell transcriptomics data, has been submitted to GEO as a private accession and is currently under embargo pending publication of a separate manuscript focused on those analyses. Access to the private dataset can be requested from the corresponding author and will be granted upon publication of the related manuscript.

## CODE AVAILABILITY

All the codes used for analysis are available at https://github.com/ojha1729/MILES#.

## ACKNOWLEDGMENTS

We thank Dr. Gregory Gibson for discussions and guidance during data analysis and interpretation. We also thank Dr. Anurendra Kumar for valuable discussions in the early phase of the analysis. Funding: This work was supported by the National Institutes of Health (R35GM131819 to S.S.) It was also supported by the Georgia Institute of Technology’s Systems Mass Spectrometry Core Facility. We thank the Marcus Foundation for sponsoring the work leading to generation of metabolomics data reported by us.

## Notes

### Competing Interest Statement

The authors have declared no competing interest.

