## Supplementary Information for "Multi-omics integration at cell type resolution uncovers gene-metabolite mechanisms underlying osteoarthritis heterogeneity"

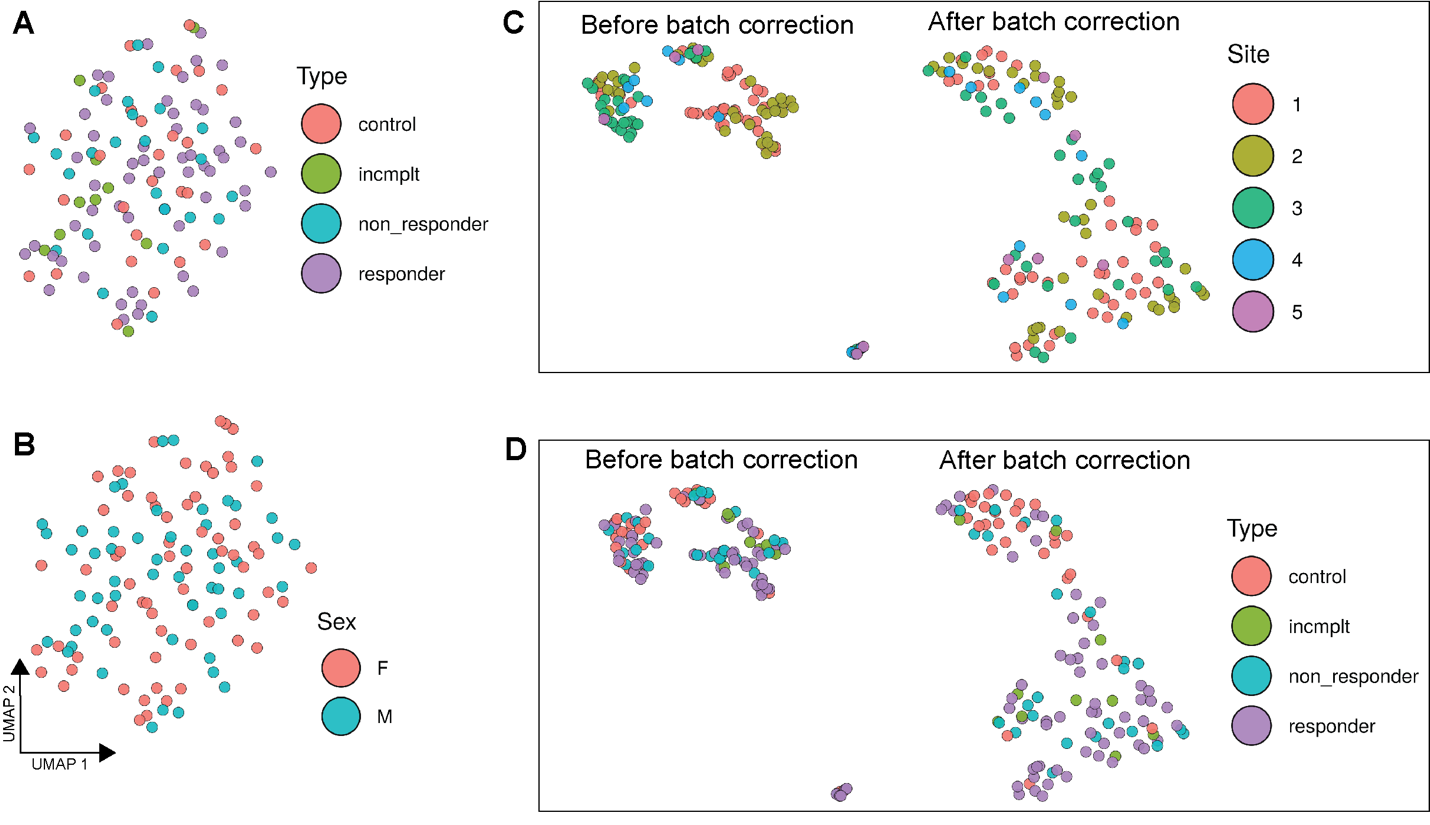

**Supplementary Figure 1:** UMAP visualizations of pseudo-bulk transcriptomics and bulk metabolomics profiles of 119 patient BMAC samples. **(A,B)** UMAP of pseudo-bulk transcriptomics profiles colored according to type of patients and sex, respectively. “Type” refers to treatment status in the cell therapy clinical trial, with the four types being “responder”, “non_responder”, “incomplete” and “control”. **(C,D)** UMAP of bulk metabolomics profiles before (left) and after (right) batch correction, colored according to site of sample collection (C) and type of patients (D), respectively.

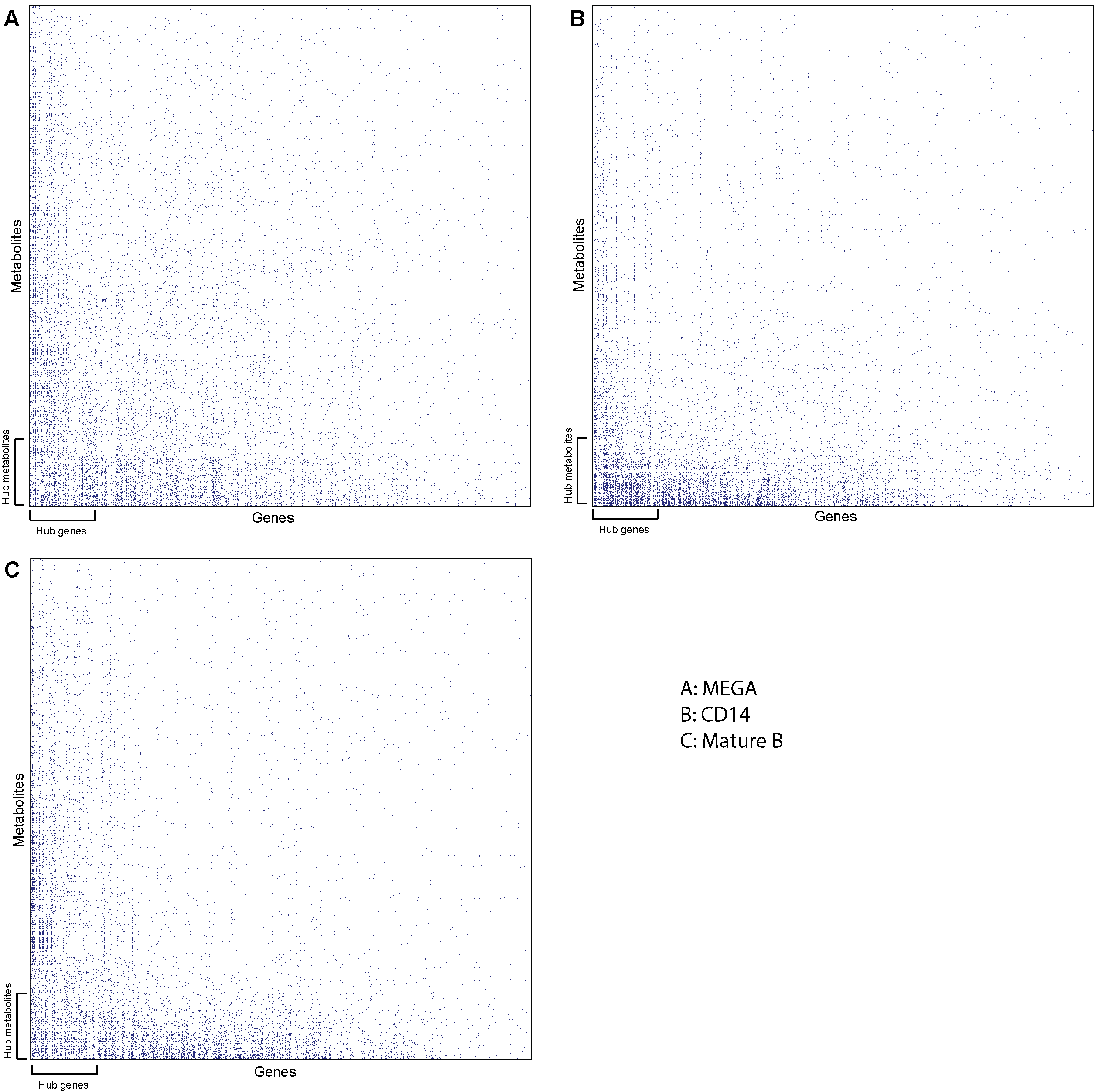

**Supplementary Figure 2:** Heatmaps highlighting associations between genes and metabolites for three select cell types. Each blue dot represents a significant association (nominal p-value < 0.005). Panels A, B, and C show results related to megakaryocytes (MEGA), CD14 Monocytes, and Mature B cells, respectively. Hub genes and metabolites are shown as leftmost columns and rows near the bottom respectively.

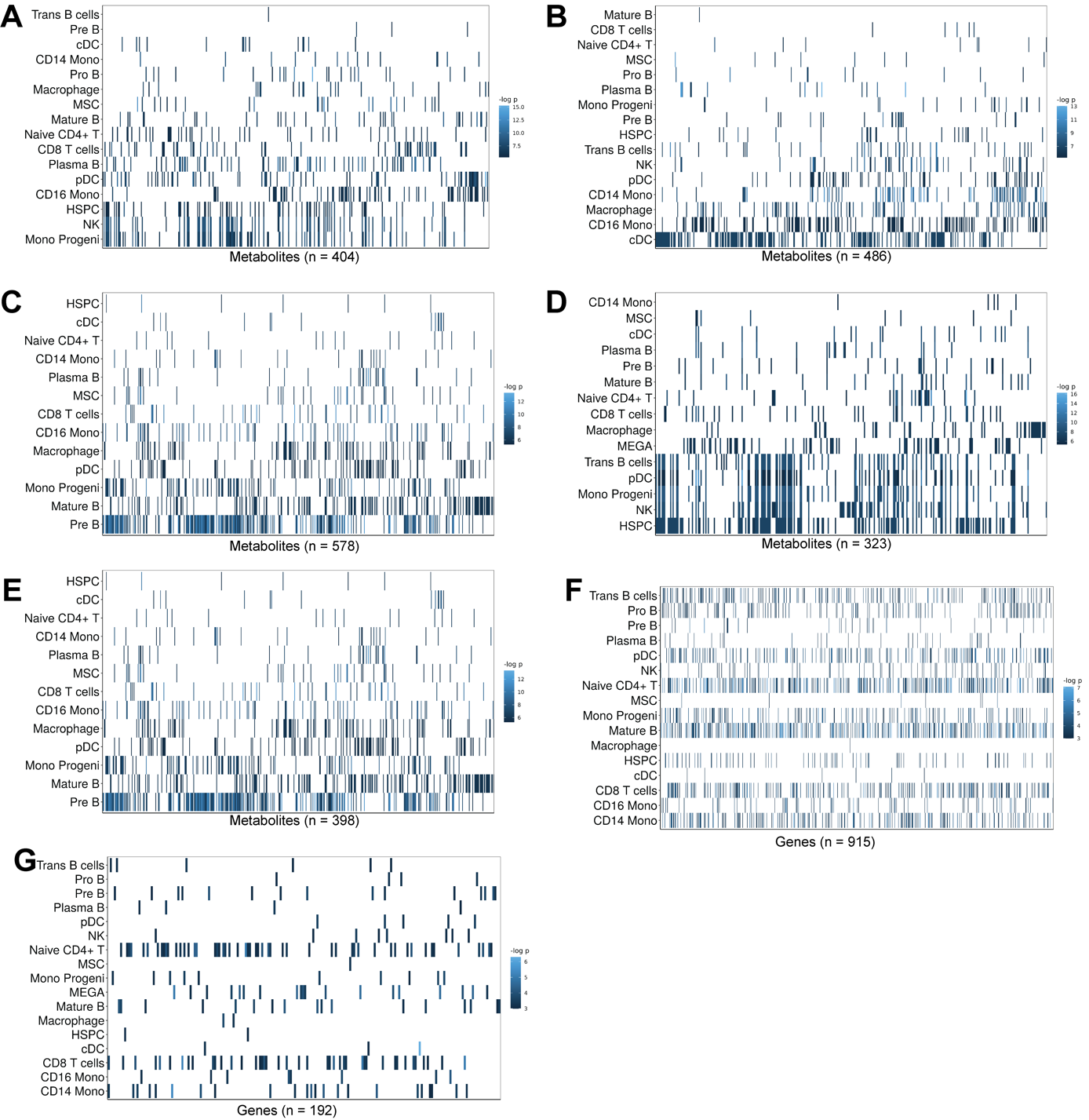

**Supplementary Figure 3:** **(A-E)** The strength of association between metabolites (columns) and pan-cell type hub genes BMP3 (A), ALPL (B), TPSB2 (C), SULF1 (D), ALDH1A3 (E) respectively, in each cell type (rows). **(F)** The strength of association between a pan-cell type hub metabolite – “SM(d42:2)” – and genes, in each cell type. **(G)** Strength of association between a cell-type specific hub metabolite, “FA(18:3)” (hub for CD8 T cells and Naïve CD4+ T cells) and genes, in each cell type.

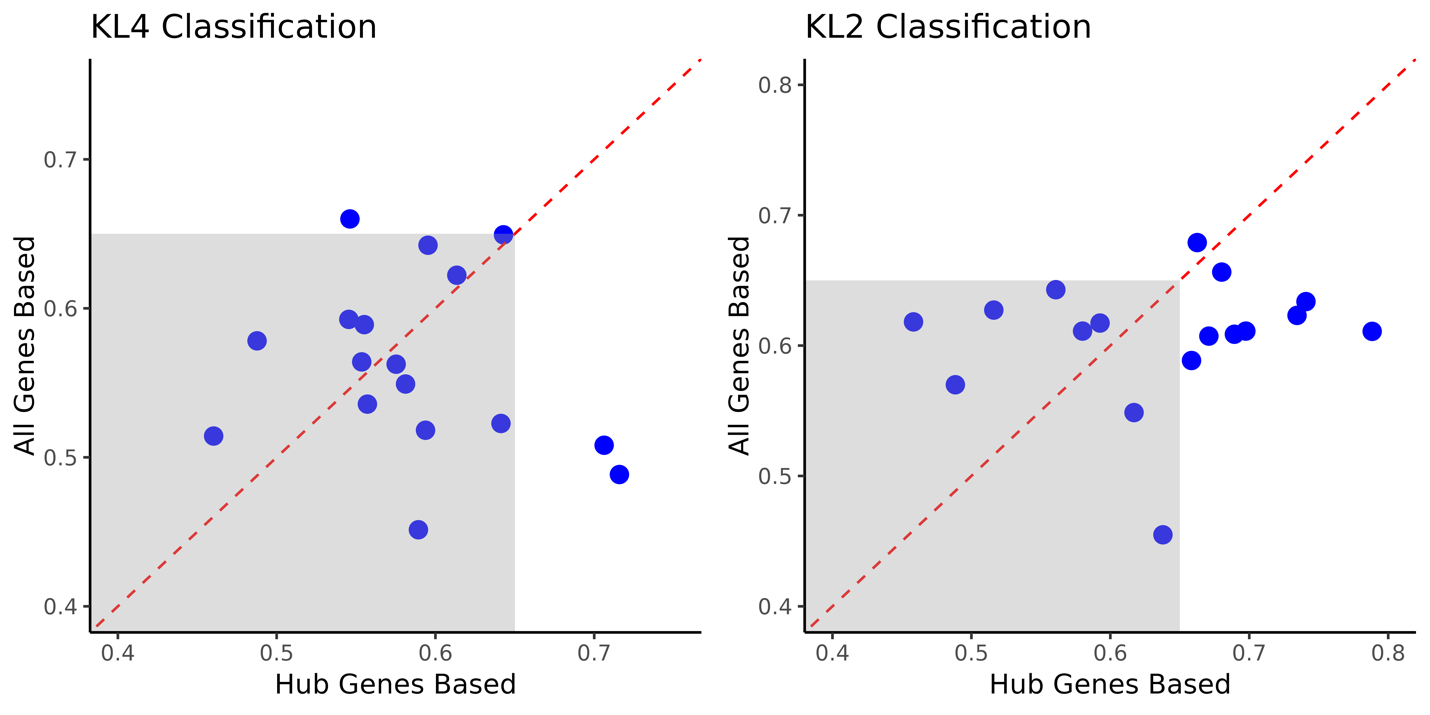

**Supplementary Figure 4:** AUROC values (on test data) for Random Forest-based prediction of KL scores (KL4 vs rest on the left and KL2 vs rest on the right) using the pseudo-bulk transcriptomic profile of each cell type. Each point represents a cell type. In each plot, the y-axis shows the values when all the genes are used for training and prediction and the x-axis shows the values when only hub genes from the covariation analysis are used. (Note that hub genes selection did not utilize the KL score information in any way.) Points outside the greyed area are cases where at least one method has a strong predictive performance (AUROC >= 0.65). In the KL2 classification, several cell types have AUROC above this threshold when using hub genes but not when using all genes. In the KL4 classification, two cell types show an AUROC > 0.7 when using hub genes and no cell type yields this level of performance when using all genes for prediction.
