## Supplementary Data Description for "Multi-omics integration at cell type resolution uncovers gene-metabolite mechanisms underlying osteoarthritis heterogeneity"

**Supplementary Data S1A, S1B:** Results for covariation analysis between each (metabolite, gene) pair in each cell type. The first column contains the metabolite names, the second column contains the gene names, the third column contains the cell-type for which the gene expression was considered, the fourth column contains the ‘p-values’ for the strength of association, and the last column contains the adjusted p-values (adjusted for multiple hypothesis tests done with each cell type). The shown triplets (metabolite, gene, cell type) in S1A are only those that have nominal p-values < 0.005. The shown triplets (metabolite, gene, cell type) in S1B are only those that have adjusted p-values < 0.05. Thus, S1B is a subset of the complete map of S1A, with more stringent criteria for inclusion.

**Supplementary Data S2:** Table containing the number of associated metabolites for each hub gene in each cell type. The first column shows the name of a hub gene (Methods), the second column contains the cell type for which the gene is considered as hub, the third column contains the number of associated metabolites (nominal p-value < 0.005), the fourth column indicates whether the given hub gene is a pan-cell-type hub gene (Methods), the fifth column indicates whether the given hub gene is a cell-type specific hub gene of the cell type in that row.

**Supplementary Data S3:** Table containing the number of associated genes for each hub metabolite in each cell type. The first column shows the name of a hub metabolite (Methods), the second column contains the cell type for which the metabolite is considered as hub, the third column contains the number of associated genes (nominal p-value < 0.005), the fourth column indicates whether the given hub metabolite is a pan-cell-type hub metabolite (Methods), the fifth column indicates whether the given hub metabolite is a cell-type specific hub metabolite of the cell type in that row.

**Supplementary Data S4:** Table showing the results related to Gene Set Enrichment Analysis of hub genes for each cell type. The first column contains the name of the broad category of the Gene Ontology (GO) term. The second column contains the cell type for which hub genes were considered. The third and fourth columns contain the GO term and GO ID, respectively. Next three columns contain the information related to the fold enrichment. The last column contains the p-value of the enrichment.

**Supplementary Data S5:** Table showing (gene, metabolite) pairs that exhibit significant association (p-value < 0.005) based on covariation analysis and belong to the same metabolic pathway. The first three columns are (metabolite, gene, cell type) triplets. The fourth column is the p-value of the strength of association between the metabolite and the gene expression in the cell type of that row. The fifth column contains the adjusted p-value at cell type level. The sixth column is the pathway code from KEGG database. The last column is the name of the common pathway to which the gene and metabolite belong.

**Supplementary Data S6:** Table showing balanced accuracy on test sets, for the prediction of metabolite abundances from the transcriptome of each cell type (Methods). The highlighted colors are those instances where test balanced accuracy is greater than 0.65. The columns titled Bulk, CT Proportion, and Confounders are the test balanced accuracies when only the bulk transcriptome, only the cell type proportion, and only the confounders were used, respectively, to predict the metabolite abundance. The column titled Count represents the number of cell types where the test balanced accuracy for the metabolite in that row is greater than 0.65. Next 17 columns are used to define cell-type-specific and pan-cell-type predictable metabolites (Methods).

**Supplementary Data S7:** Table containing the Area Under ROC curve (AUROC or AUC) for the trained random forest classifier of KL group (KL-X vs Rest, X=2 and 4) using the transcriptome of each cell-type (Methods). Evaluations are from test data (under 70:30 train-test split). The best three cell-types in terms of AUC are highlighted in colors (orange for KL-2 vs Rest classification and green for KL-4 vs Rest classification).

**Supplementary Data S8:** Table containing the list of metabolites that are significantly more abundant in KL-X (X= 2 and 4) when compared against rest based on a proportion test (see Methods). The last two columns contain the p-values based on the proportion test.

**Supplementary Data S9:** Table containing the set of metabolites that are significantly more abundant in KL-X group (X=2 and 4) and are predictable using transcriptome of any (or multiple) cell types. The table contains the p-values based of the differential abundance test and the test accuracies of prediction using cell-type transcriptome profile.

**Supplementary Data S10:** Table containing the cluster scores for interconnection between hub genes and hub metabolites in each cell-type. The first column contains the names of the cell-types. The second column contains the cluster score calculated using the NetworkX library in Python. The last column contains the empirical p-values for the cluster scores (Methods).

**Supplementary Data S11:** Table containing the degree (the number of associated metabolite) for each gene in each cell-type. The column named ‘Sum’ contains the total degree of each gene across all cell-types. The next columns contain the difference between the degree of a gene in a cell-type and average of its degree across all the other cell-types.
